# Graphene microelectrode arrays, 4D structured illumination microscopy, and a machine learning-based spike sorting algorithm permit the analysis of ultrastructural neuronal changes during neuronal signalling in a model of Niemann-Pick disease type C

**DOI:** 10.1101/2024.02.22.581570

**Authors:** Meng Lu, Ernestine Hui, Marius Brockhoff, Jakob Trauble, Ana Fernandez-Villegas, Oliver J Burton, Jacob Lamb, Edward Ward, Philippa J Hooper, Wadood Tadbier, Nino F Laubli, Stephan Hofmann, Clemens F Kaminski, Antonio Lombardo, Gabriele S Kaminski Schierle

## Abstract

Simultaneously recording network activity and ultrastructural changes of the synapse is essential for advancing our understanding of the basis of neuronal functions. However, the rapid millisecond-scale fluctuations in neuronal activity and the subtle sub-diffraction resolution changes of synaptic morphology pose significant challenges to this endeavour. Here, we use graphene microelectrode arrays (G-MEAs) to address these challenges, as they are compatible with high spatial resolution imaging across various scales as well as high temporal resolution electrophysiological recordings. Furthermore, alongside G-MEAs, we deploy an easy-to-implement machine learning-based algorithm to efficiently process the large datasets collected from MEA recordings. We demonstrate that the combined use of G-MEAs, machine learning (ML)-based spike analysis, and four-dimensional (4D) structured illumination microscopy (SIM) enables the monitoring of the impact of disease progression on hippocampal neurons which have been treated with an intracellular cholesterol transport inhibitor mimicking Niemann-Pick disease type C (NPC) and show that synaptic boutons, compared to untreated controls, significantly increase in size, which leads to a loss in neuronal signalling capacity.

## Introduction

The advent of microelectrode arrays (MEAs) has facilitated long-term, large-scale monitoring of local field potentials of neurons, offering non-invasive recordings of a broad range of spatial and temporal neuronal signals, thus, surpassing traditional patch-clamp recordings^1–4^. Nevertheless, non-transparent MEAs often face limitations in fundamental and therapeutic neuroscience research, owing to their incompatibility with advanced imaging setups as well as software constraints required for spike analysis.

To address these problems in part, transparent MEAs have been under development in the recent decade to allow for the combination of optical microscopy and electrophysiology^5–9^. However, the design of transparent MEAs often comes with a trade-off between the low impedance values of individual electrodes, required to achieve high signal-to-noise ratios in the electrophysiological recordings, and the transparency of the imaging field of view (FOV) required for optical microscopy. State-of-the-art commercial indium tin oxide (ITO) MEAs display impedance values in the range of around 250 kΩ at 1 kHz, which is sufficient to detect neuronal activity and offer electrode transparency of about 80 % in the visible light spectrum^7,8,10,11^. However, despite the improved transparency, the electrodes themselves are still highly visible under the microscope, which leads to the uneven attenuation of the excitation light and, by that, to distortions, such as the warping of excitation patterns, resulting in artefacts of reconstructed images obtained from high-resolution imaging systems such as structured illumination microscopy (SIM).

Graphene, *i.e.*, a monolayer of carbon atoms tightly arranged in a honeycomb structure^12^, exhibits good electrical conductivity^13^, which is also frequency-independent up to microwave frequencies^14^, making it suitable for the detection of spontaneous neuronal activity^15,16^. Furthermore, its biocompatibility permits long-term cultivation of cells^15,17–21^, which is another requisite for long-term recordings of neuronal signals. However, most importantly, a single layer of graphene has a transparency of 97.7 % in the visible light spectrum, with a negligible reflectance of less than 0.1 %^22^, which is, thus far, the highest transparency measured amongst all electrode materials used for transparent MEAs. Therefore, graphene is an ideal material to be used in MEAs to permit the concomitant analysis of neuronal structures and signals both during *in vitro* and *in vivo* studies^23,24^. However, despite these advances in proof of principle studies, only a few publications have emerged since, which suggests that improvement is needed to make this technology applicable and accessible for addressing important biological questions.

One related limitation is that recordings taken at an electrode often include signals from multiple overlapping neurons and other electrical noise sources. Therefore, spike sorting, *i.e*., the task of distinguishing neuronal activity from background noise as well as assigning neuronal activity to its respective source neurons, is crucial to ensure proper analysis^25–27^. Recently, the application of data-driven, machine learning (ML)-based approaches to spike sorting has been proposed^28–30^. Accordingly, here, we benchmark several deep clustering methods and evaluate spike sorting performance using a large-scale, simulated dataset, before compiling these analysis tools into an easy-to-use Python script which is available as an open-source tool. As part of the open-source tool, we also provide a Python script for the analysis of fluorescence-based calcium imaging data, where the fluorescent calcium traces are bleach-corrected and calcium spikes are extracted, therefore enabling the cross-comparison of fluorescent calcium-based imaging and electrophysiological data.

We demonstrate with the combined use of graphene MEAs (G-MEAs), four-dimensional (4D) SIM and our analysis software, that we can monitor the activity and morphology of neurons across scales, *i.e.*, from the network level to the sub-neuronal domain. We particularly focus on synchronicity disruptions and morphological defects in hippocampal neurons upon lysosomal cholesterol accumulation, a phenotype mimicking Niemann-Pick disease type C (NPC). The corresponding long-term recordings over large FOVs reveal the degeneration of neuronal networks, leading to the loss of neuronal activity which is accompanied by structural changes at the single synapse level as revealed during neuronal firing.

## Results and Discussion

### G-MEAs and machine-learning-based data analysis enable the concomitant and detailed investigation of neuronal structures and signals

Our G-MEAs are designed to be compatible with commercially available recording hardware. Nevertheless, the introduced approach and techniques, including the device fabrication, offer high flexibility, such that the design can also easily be adapted for custom setups. The G-MEAs presented here enable the detection and recording of neuronal action potentials (Materials and Methods), however, there is still room to improve the fabrication protocol for G-MEAs in the future, *e.g.*, by increasing the contact between the neurons and the graphene surface in order to increase the number of electrodes that record electrical signals per batch produced^15^.

**Fig. 1a** summarises the experimental pipeline used to facilitate the applicability of G-MEAs for the study of complex neuronal degeneration over time. Primary hippocampal neurons are cultured on G-MEAs, where they are maintained for more than 21 days before optical as well as electrophysiological data are acquired and, subsequently, analysed using a spike sorting algorithm. Additionally, on day four *in vitro* (DIV4), the hippocampal neurons are transduced with adeno-associated viruses (AAV) to express the fluorescence-based calcium sensor GCaMP7b, allowing for image-based calcium activity tracking of both single neurons and neuronal networks (**Fig. 1b**).

**Figure 1.**
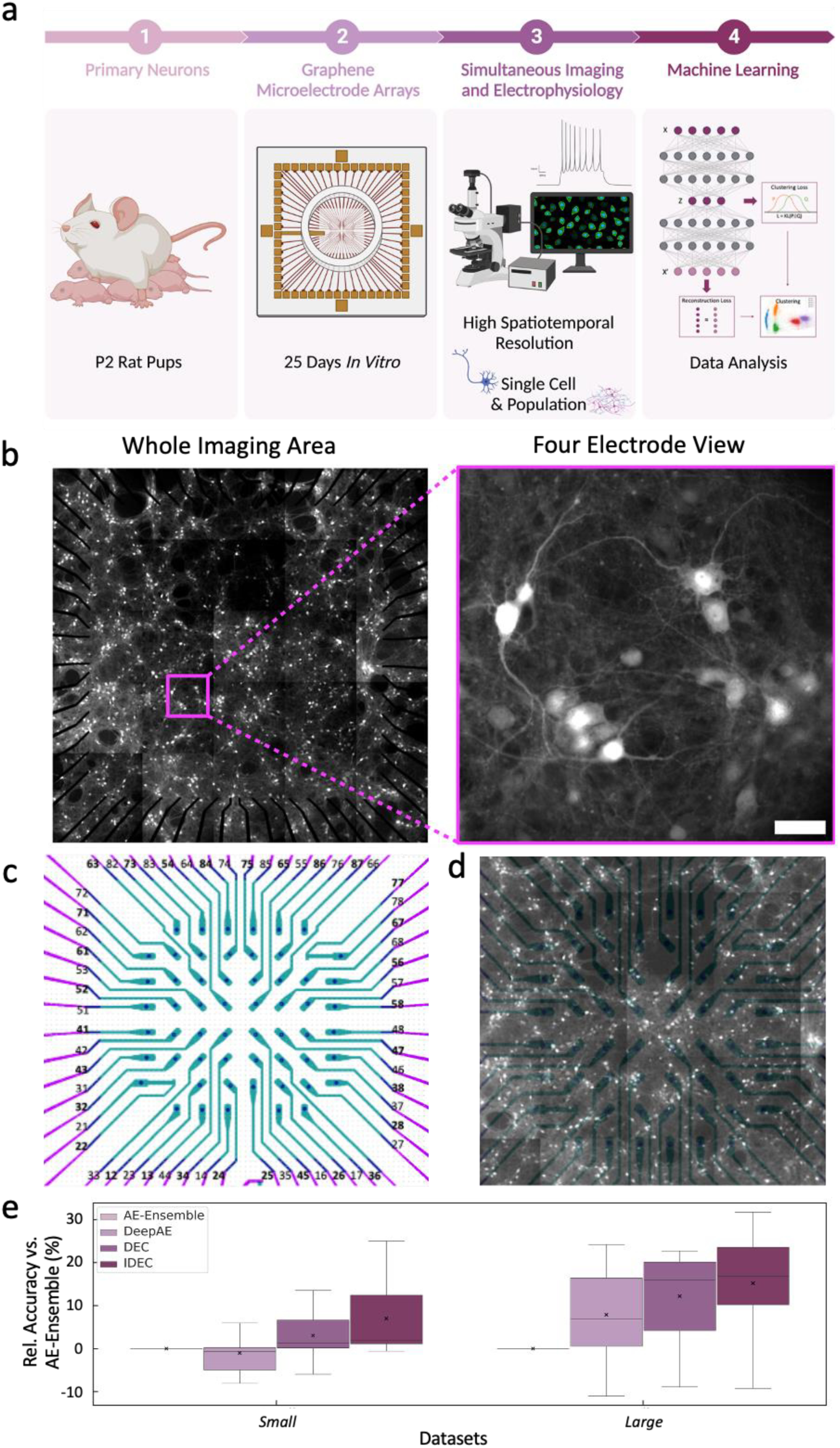
G-MEAs and ML-based data analysis enable the advanced study of neuronal structures and signals. (**a**) Experimental pipeline for the detailed investigation of neuron-related diseases using G-MEAs, consisting of post-natal day 2 (P2) hippocampal neurons being plated on transparent G-MEAs. Simultaneous electrophysiological and imaging recordings of the neurons were obtained and analysed using ML. (**b**) Left: Image showing the entire transparent area of the G-MEA with a size of 4.84 mm^2^. Right: Image highlighting a portion of the device containing four graphene microelectrodes. Scale bar: 50 µm. (**c**) An overview of the array displaying the microelectrodes and their respective numerical labels. The gold electrodes are shown in magenta and the graphene microelectrodes are shown in cyan. The reference electrode is situated between electrodes 24 and 25. (**d**) The overlay of the graphene microelectrodes’ blueprint with the image of the whole imaging area ensures correct interpretation and correlation of the orthogonally collected data. The device map is then split into quadrants to isolate each area for separate imaging techniques and to minimise phototoxic effects. A neuron located close to an electrode is then selected within each quadrant for further investigation. (**e**) Shown are classification accuracies of proposed deep clustering approaches (DeepAE, DEC, and IDEC) as well as the existing state-of-the-art AE-Ensemble approach for different simulated datasets. Marked are median (grey line), mean (black cross) and minimum/maximum (grey whiskers) accuracy. All models have been benchmarked on a large-scale simulated benchmarking dataset, containing 10 sets of data each for the *Small* and *Large* (**SI Table 3**) datasets, and results are normalised to AE-Ensemble for improved accessibility. Fig. 1a is created using Biorender.

The subsequently applied analysis software aims to extract calcium spiking rates of, or spiking synchronicity between, individual neurons. For the spike sorting task, we build on insights gained from existing models including Deep Embedding for Clustering^31^ (DEC) and Improved Deep Embedding for Clustering^32^ (IDEC). Further, the synthetic datasets currently used in the field only provide limited numbers of spike recordings (<5,000) and from a low number of source neurons (typically three)^25^, which makes them too small and simple to capture the analysis-related challenges associated with real-time recorded MEA data, we benchmark the different spike sorting algorithms using newly simulated datasets. The *Small* and *Large* datasets (**SI Tables 1** and **2**) presented here have been prepared *via* NeuroCube^33^ and consist of 10 sets of spike recordings from five source neurons containing around 100,000 (small) and 1,100,000 (large) spikes each, respectively.

Both DEC and IDEC result in improved spike sorting accuracy compared to the state-of-the-art Autoencoder-Ensemble^28^ (AE-Ensemble) as well as a deep autoencoder approach (Deep AE) which forms the basis of both DEC and IDEC (**Fig. 1e**). Further details on how the individual models are trained are provided in **Materials and Methods**. Across the *Small* datasets, DEC and IDEC, on average, show a relative improvement in accuracy of 3.03 ± 11.32 % (mean ± standard deviation) and 7.00 ± 8.87 % if compared to the AE-Ensemble, respectively. For *Large* datasets, the relative improvement of DEC and IDEC increases to 12.20 ± 10.65 % and 15.21 ± 12.26 %, respectively, while DeepAE shows a relative change of −1.02 ± 5.48 % for the *Small* and 7.89 ± 11.13 % for the *Large* datasets if compared to AE-Ensemble.

The outperformance of DEC and IDEC compared to AE-Ensemble with increasing dataset sizes may be based on DEC and IDEC possessing more trainable parameters and, therefore, being able to extract spike features from big datasets more efficiently. This would suggest that their performance scales positively with training dataset size, which is crucial for future electrophysiological applications^30^. However, a performance improvement caused by the optimised training methods cannot be excluded. Additionally, although DeepAE has the same number of trainable parameters as DEC and IDEC, the joint clustering and feature extraction, which is only used during the training of DEC and IDEC, may improve their performance.

Following the benchmarking of the ML-based method, we have applied our approach for the analysis of the correlative network activity of hippocampal neurons grown on G-MEAs. As illustrated in **Fig. 2**, G-MEAs permit the independent as well as simultaneous acquisition of electrophysiological and imaging data, with **Fig. 2a** displaying the recording of typical neuronal spike shapes. The firing behaviour observed in our study, including burst-like firing (**Fig. 2b**, left) and single spike firing events (**Fig. 2b**, right), aligns with earlier recordings from hippocampal neurons^34^. Additionally, building on the G-MEA’s high transparency, the transparent G-MEAs facilitate the investigation of spontaneous calcium release, an important phenomenon not directly coupled to the firing of an action potential^35^.

**Figure 2.**
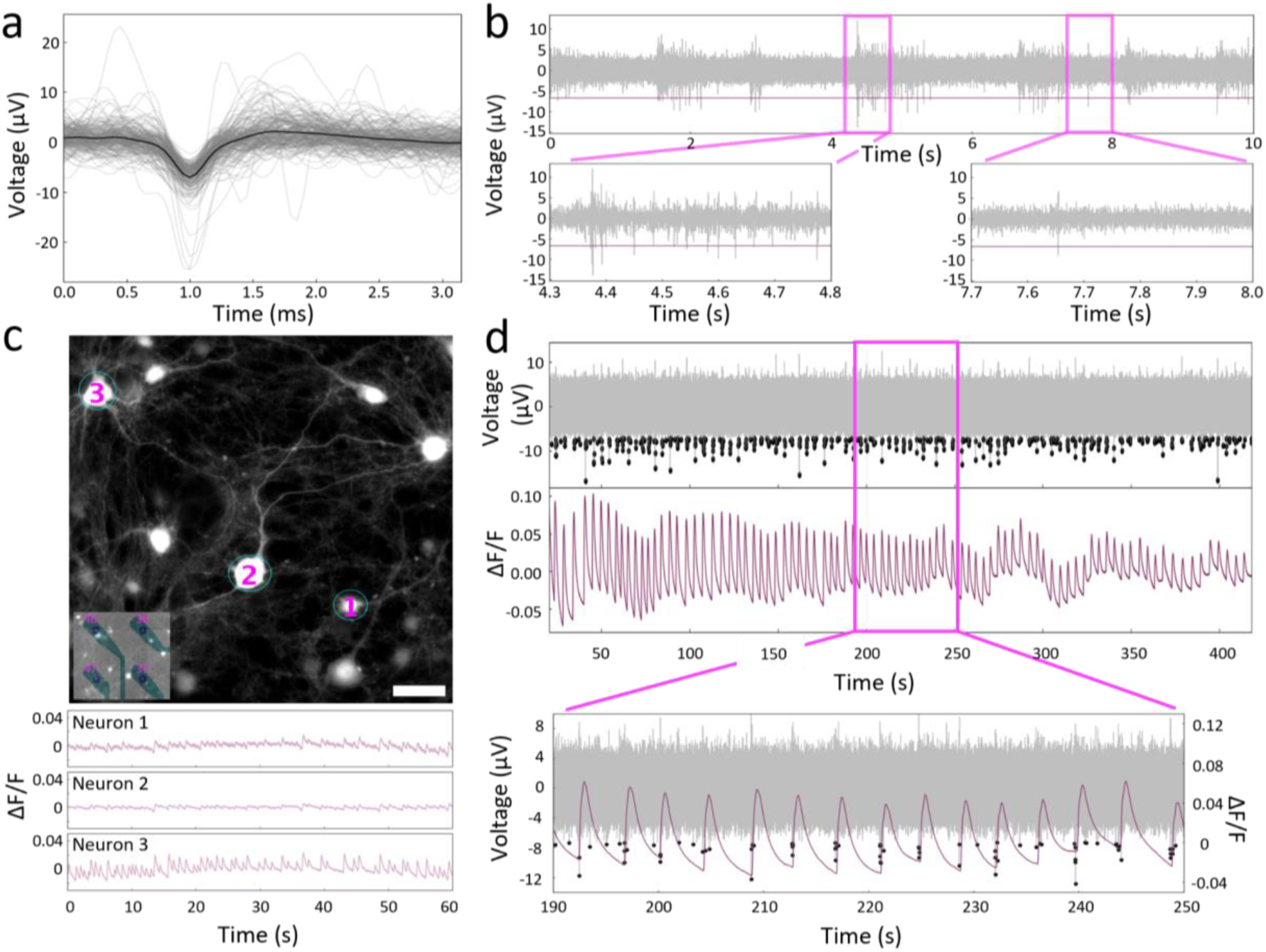
G-MEAs enable simultaneous imaging and electrophysiology. (**a**) Neuronal spike shapes recorded by G-MEAs. The bold line indicates the average spike shape. *N*=200. (**b**) Filtered recording acquired by a single electrode. Enlarged regions illustrate different patterns of hippocampal neuronal activity, *i.e.*, burst (left) and single spike firing events (right). The horizontal, magenta line indicates the threshold for spike detection. (**c**) Calcium imaging of mature hippocampal neurons on G-MEAs with the respective positions of electrodes indicated in the lower left corner. Bottom: Change in amplitude over basal fluorescence intensity (ΔF/F) over time for the three neurons (DIV 24) labelled as 1-3 (top). Scale bar: 50 µm. (**d**) G-MEAs enable simultaneous acquisition of both local electrophysiological and calcium recordings as illustrated by the co-occurring spiking events. Simultaneously acquired electrophysiological recordings (grey, left y-axis) including detected spikes (black dots) are manually aligned with the respective calcium signals (magenta, right y-axis) of a neuron. The imaged neuron was located close to the corresponding electrode.

**Fig. 2c** presents a fluorescence microscopy image from an area containing four graphene microelectrodes, as illustrated in the insert, and (**Fig. 2d**) shows the simultaneous calcium imaging and electrophysiological recordings of neurons that are close to the electrodes. These neurons display coherent co-occurrence of spiking events in both signal recording domains. Additionally, in contrast to previous studies that combine MEAs with simultaneous imaging to record neuronal activity^15–19,22–24,36–38^, our transparent MEAs offer the unique advantage of enabling imaging directly above the electrodes in a locally confined area. This feature allows for precise correlation of the neuronal activity of a single neuron at a specific electrode with its corresponding calcium spike, as demonstrated in **Fig. 2d**, providing the capability to identify defects that may impact either the neuron’s ability to fire an action potential or its ability to release neurotransmitters. For example, correlated recordings obtained from different locations (**Supplementary Fig. 1**) can expose significant variations in behaviour, such as reduced matching accuracy between the two signal domains or varying firing rates, which may be indicative of early disease. Further, local correlation can be of relevance for future applications, such as for optogenetic- or electric-based localised stimulation and the investigation of signal transmission across neuronal networks. Additionally, this correlation is relevant for the testing of various neuronal drugs, where it is crucial to determine the pathway a drug affects. For instance, calcium channel blockers, some of which are currently undergoing clinical trials for treating neurodegenerative diseases and other brain-related disorders, such as stroke^39^, exert their effects by solely influencing intracellular calcium influx, thereby blocking neurotransmitter release. However, certain drugs may exclusively modulate the neuron’s capacity to fire an action potential. Hence, understanding these distinctions is vital for optimising long-term treatment strategies for patients with neurodegenerative diseases.

### Quantitative analysis of the cellular Niemann-Pick disease type C phenotype reveals a loss of synchronicity in neuronal firing patterns and structural impairments of primary hippocampal neurons

After validating the G-MEA’s suitability to permit simultaneous electrophysiology and optical recordings, as well as verifying the performance of our ML-based analysis, we determine the effect of the intracellular cholesterol transport inhibitor U18666A, which mimics Niemann-Pick disease type C (NPC), on primary hippocampal neurons. On a global level, we observe that U18666A treatment induces substantial neuronal degeneration over time. To quantify disease progression, both individual and simultaneous readouts of neuronal signals and imaging data are captured (**Fig. 3**). The electrophysiological and the calcium imaging data reveal that neurons treated with U18666A display a significant decline in spike rate and synchronicity after four days when compared to control conditions (**Fig. 3a**), suggesting that U18666A strongly affects neuronal networks. Interestingly, while the electrophysiological activity starts to decline shortly after treatment, the calcium spike rate in the U18666A condition first exhibits an increase before declining on the subsequent days, suggesting that U18666A treatment first impacts calcium homeostasis as shown recently for models of NPC^40^. Furthermore, the detected loss in activity for U18666A treated cells across the different signal types, as presented in **Fig. 3a**, is further accompanied, and potentially enhanced, by a decrease in the overall cell density (**Fig. 3b**), similarly to what has been observed for NPC patient-specific induced pluripotent stem cells (iPSCs)^41^.

**Figure 3.**
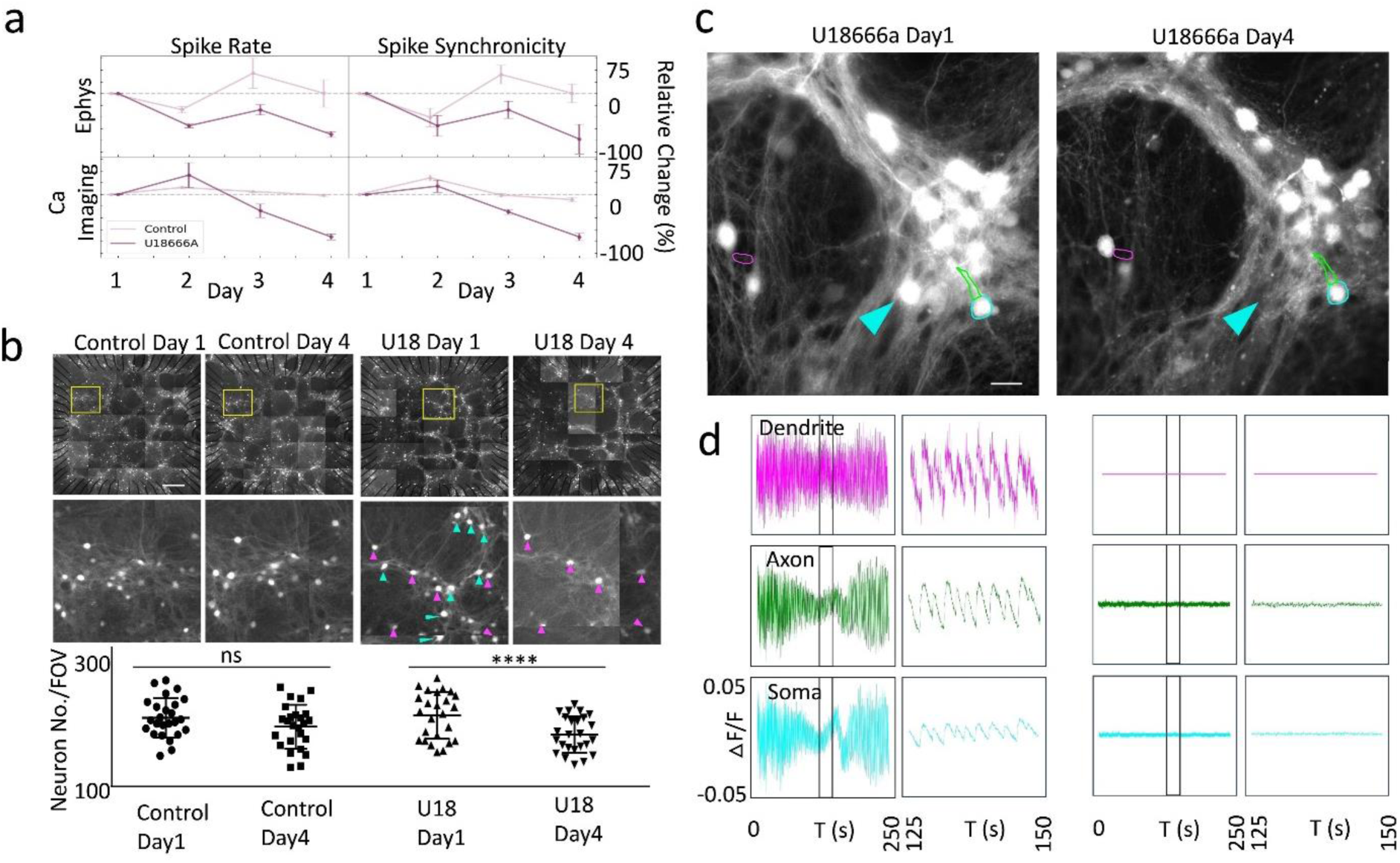
Capturing the deterioration in neuronal structures and activity in a Niemann-Pick disease type C model from a population to a sub-neuronal level. (**a**) The effects of U18666A on the electrophysiological (upper row) and calcium imaging activity (lower row) of primary hippocampal neurons over four days, normalised by the activity on day 1 (control in light purple; U18666A in dark purple). Shown is a relative change for the spike rate (left column) and spike synchronicity (measured as mutual information^42^, right column). Error bars indicate the standard error of means. (**b**) Calcium imaging of the neuronal population across the whole FOV (top panel) and a single FOV (middle panel) in both control (left group) and U18666A-treated samples (right group). Left panel: samples on day one (DIV 24) before U18666A treatment. Right panel: samples on day four (DIV 27) after U18666A treatment. The U18666A treatment, finally, resulted in a loss of neurons. The cyan arrows highlight the neurons that have disappeared during U18666A treatment, and the magenta arrows highlight the surviving neurons. Scale bars: 100 µm for whole FOV, 20 µm for enlarged FOV. U18: U18666A. Bottom panel: Quantification of neuron number per FOV of device for U18666A treatment *vs.* control. Twenty-five images per FOV of a device were acquired in each experiment. Four independent experiments were performed. (**c**) Neuronal structures imaged by widefield microscopy. Sub-neuronal structures highlighted in the figure are a dendrite (magenta), an axon (green), and a soma (cyan). A neuron that disappeared after the four-day treatment is highlighted by a cyan arrow. (**d**) Changes in the amplitude of the fluorescent calcium sensor extracted from the above neuronal structures are depicted over a time window of 250 s. Y-axis: ΔF/F with a scale of −0.05 to 0.05 for all plots. The short time window (25 s) is indicated by black rectangles in the full-time window panel on the left.

To investigate the potential cellular mechanisms causing the above-observed defects in further detail, individual sub-neuronal structures, namely somas, axons, and dendrites, are investigated using widefield microscopy. The corresponding analysis reveals substantial changes in the amplitude of the calcium activity over time (**Fig. 3c and d**). Specifically, the four-day treatment with U18666A leads a loss of neuronal structures, such as shown in **Fig. 3c**, which is accompanied by the depletion of calcium activity in these structures (**Fig. 3d**). In contrast, control neurons display clear and regular calcium spikes that are consistently present throughout the whole recording (**SI Fig. 2**). Observations in cells treated with U18666A are in alignment with earlier *in vivo* and *in vitro* experiments that have consistently identified synaptic plasticity defects in NPC models, such as arising from either mutation in the NPC1 gene or inhibition of NPC1 by U18666A. Reported defects manifest themselves as increased synaptic cholesterol levels^43^, impaired Long-Term Potentiation (LTP)^44^, the formation of axonal spheroids^45^, reduced dendritic spine density^46^, and ultimately, neuronal loss^40,47^ — the latter two of which are also observed in our model system (**Fig. 3b-d**). Nevertheless, the distinct advantage of our study lies in its capability to not only unveil defects in neuronal activity but to further allow for the precise and simultaneous identification of the corresponding degenerating neuronal structures.

### SIM imaging reveals that sub-neuronal structures dynamically alter their shape during calcium signalling

Leveraging the superior transparency of G-MEAs, our MEAs are combined with high-resolution microscopy, more specifically structured illumination microscopy (SIM), to investigate the underlying mechanisms of sub-neuronal structural changes in further detail. When compared to widefield microscopy^48^, SIM imaging offers a twofold gain in resolution, providing highly resolved sub-neuronal structures that can be quantitatively analysed to obtain information about morphology, size, and activity. Accordingly, this approach enables us to observe the dynamic nature of these sub-neuronal structures, tracing intricate morphological changes over time as well as to correlate them with the underlying calcium activity (**Fig. 4**). We, therefore, capture a time-lapse sequence of three-dimensional (3D) sub-neuronal structures (**Fig. 4a**) on DIV 27, *i.e.*, 23 days after transduction with the calcium marker GCaMP7b, before reconstructing the images to obtain a 4D stack containing spatial-temporal information of calcium activity at sub-neuronal levels (**Supplementary Movie 1 and Fig. 4b**). As shown in the projection view of the 4D stack in **Fig. 4b**, the neuronal ultrastructures can be captured and analysed in the reconstructed data, which demonstrates a significant improvement in revealing the structural details of live neurons compared to the widefield microscopy counterpart.

**Figure 4.**
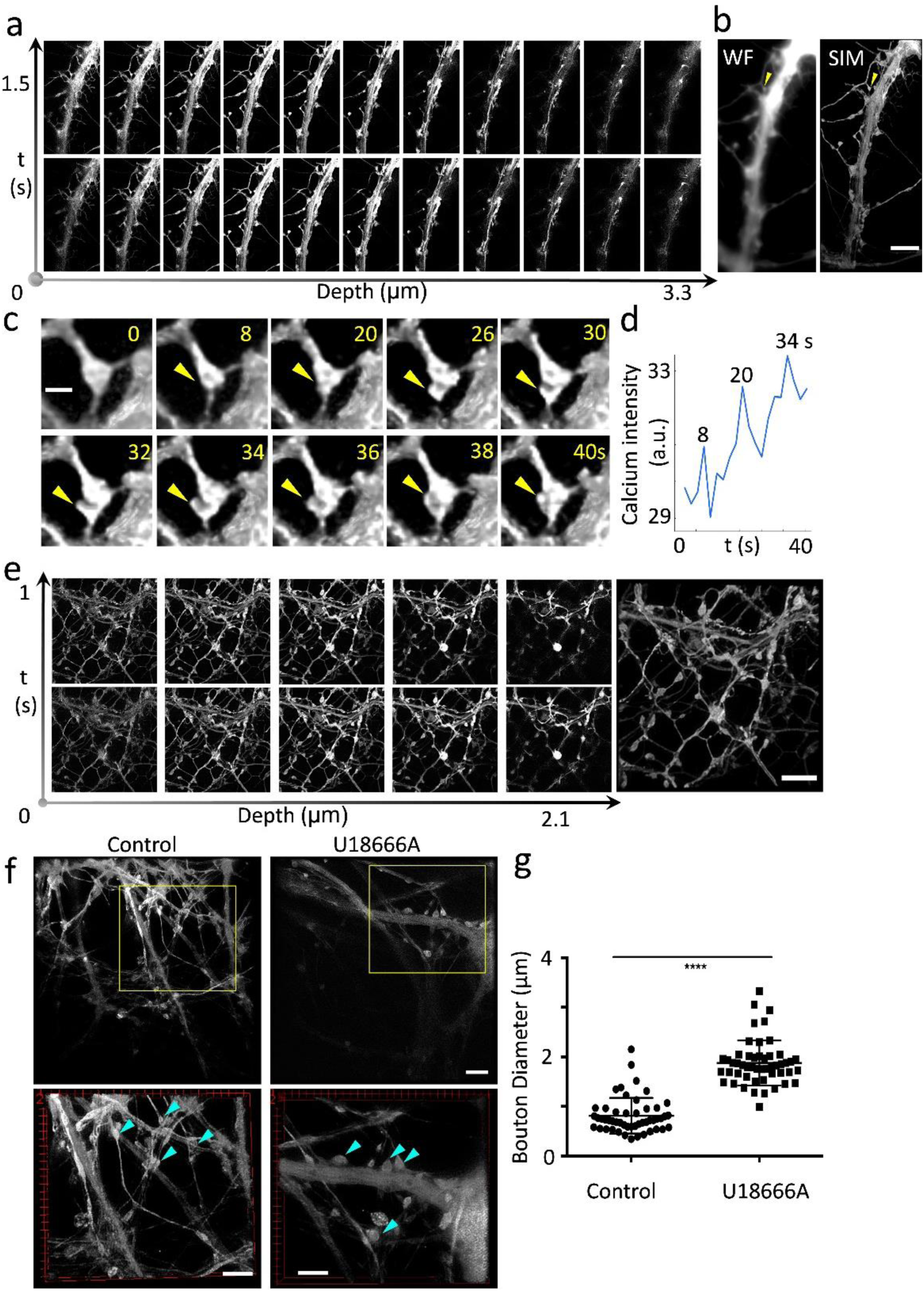
Four-dimensional SIM reveals that neuronal ultrastructures can dynamically alter their shape during calcium signalling and due to stress induced by U18666A. (**a**) A representative neuronal structure is imaged using sectioning SIM in a time-lapse sequence. (**b**) A projection view of the 4D data reconstructed from the entire series of images to illustrate changes in structure and calcium activity over time. A 4D widefield (WF) version of the reconstructed images is displayed as a comparison. The yellow arrow points to a neuronal ultrastructure that undergoes significant structural changes over a timeframe of 40 s (Supplementary Movie 1 and, enlarged, in (c)). Scale bar: 5 μm. (**c**) The enlarged view of the neuronal ultrastructure highlighted in (b) shows the structural change during calcium activity. The yellow arrows indicate the local enlargement of the neuronal ultrastructure. The time in seconds is indicated in the top right corner. Scale bar: 0.5 μm. (**d**) The calcium intensity data extracted from the image sequence of the neuronal ultrastructure is shown in (c). The spikes are labelled with the corresponding time stamps. (**e**) A complex neuronal network was captured using sectioning SIM in a time-lapse sequence. Scale bar: 5 μm. (**f**) Representative SIM images showing control neurons and neurons treated with U18666A. Top panel: whole FOV from SIM images. Bottom panel: enlarged view from the yellow boxed regions above. Neuronal ultrastructures are highlighted with cyan arrows. Scale bars: 5 μm. (**g**) Quantitative analysis of the diameter of synaptic boutons reveals a significant increase following U18666A treatment. *N*=50. Mean ± SD. \*\*\*\**P* < 0.0001. Statistical significance was evaluated using the Student’s t-test of 6 independent experimental repeats.

Using 4D SIM imaging data, we then analyse how the synaptic structure changes during neuronal activity. **Fig. 4c** presents an enlarged view of a neuronal ultrastructure highlighted by a yellow arrow in **Fig. 4b**, with the calcium intensity data extracted from the imaging data being shown in **Fig. 4d**. The graph showcases the temporal relationship between structural modifications and calcium activity or intensity, with the local calcium fluorescence analysis (**Fig. 4d**) exposing three calcium peaks in a time window of 40 seconds which coincide with a reformation of the synapse. Transient enlargements of neuronal ultrastructures, such as dendritic spines and synaptic boutons during neuronal activity have been observed previously in hippocampal neurons, which might be caused by the underlying morphological changes we observe during calcium spike events^49^. However, whether they are triggered by calcium influx or not needs to be further investigated.

Next, we apply SIM to elucidate the details of the complex network formed by hippocampal neurons, as depicted in **Fig. 4e** and **Supplementary Movie 2**, with a specific focus on neuronal ultrastructures. The axons and dendrites intricately interweave, forming a dense mesh that signifies a high degree of interconnectivity crucial for efficient signal transmission and neural plasticity. This interconnected network plays a pivotal role in supporting spontaneous activity in hippocampal neurons as also demonstrated by the distinct multi-layered axonal and dendritic projections revealed in the sectional images, with dendritic spines and synaptic boutons being resolved at different focal planes.

Finally, we apply 4D SIM to investigate the sub-neuronal structural changes following treatment with U18666A (**Supplementary Movie 3 and 4**). **Fig. 4f** shows representative reconstructed images of both conditions, with neuronal ultrastructures highlighted using cyan arrows. Quantitative analysis of synaptic boutons after U18666A treatment reveal a significant increase in the size after U18666A treatment (**Fig. 4g**). Consequently, the neuronal defects at the network level (**Fig. 3**) can likely be attributed to the presence of dysfunctional synapses, characterised by activity loss, enlarged structures, and even the loss of synapses themselves (**Fig. 4f-g**). While these observations of neuronal deteriorations are in agreement with previous studies reporting on the progression of NPC^50^, thus far, investigations into structural defects at the sub-neuronal level, particularly at individual synapses, *i.e*., the primary loci of structural plasticity, have remained largely unexplored. Hence, in our investigations, we advance the field by employing 4D SIM to permit the spontaneous examination of individual neuronal ultrastructures and their correlated structural changes for the first time. Further, the observed increase in synaptic bouton size, indicative of synaptic dysfunction, aligns with the synaptic theory of neurodegeneration^51^, which postulates that synaptic deficits precede neuronal cell death and are pivotal drivers of disease progression.

As presented here, the 4D SIM data recorded on our G-MEAs are capable of providing direct insights into how sub-neuronal structures change during neuronal activity, shedding light on the intricacies of structural plasticity at the individual synapse level. This offers a more profound understanding of how neurodegeneration and synaptic loss may unfold, and impact calcium homeostasis but also other organelles^46^. Indeed, in a recent study, we have demonstrated that U18666A significantly impairs the tubular endoplasmic reticulum (ER) in COS-7 cells using SIM^52,53^. Such impairment in the tubular ER can be particularly detrimental if observed in neuronal cells, as the axon primarily consists of tubular ER. Additionally, given that synapses receive crucial components such as proteins, lipids, and calcium *via* the tubular ER, damage to the tubular ER in the axon may significantly impact synaptic function. Therefore, future work may aim to combine electrophysiology with a SIM analysis of the ER to experimentally test this hypothesis.

### Conclusion

The simultaneous use of orthogonal investigation techniques is essential to further our understanding of complex molecular processes, such as those involved in the progression of neurodegenerative diseases. Here, we build on the unique properties offered by graphene, namely its conductivity and transparency, to permit the optical as well as electrophysiological study of neuronal cells across scales. By pairing the device with machine learning-centred analysis algorithms, we evaluate its suitability for the image-based recording of calcium spikes. We further investigate the effects of U18666A, mimicking Niemann-Pick’s disease type C, where a loss of network synchronicity, as well as individual neurons, are detected, before analysing the sub-neuronal features in greater detail through SIM, revealing changes in the morphology of synapses in treated neurons. Our methodology, which integrates ML-based spike analysis with simultaneous electrophysiology and fluorescence imaging-based calcium activity measurements, will not only be highly beneficial for the study of synaptic dysfunction/loss/increase in models of neurodegenerative diseases but also in models of schizophrenia and epilepsy, respectively. However, to make our methodology more available, some of the technical challenges associated with the fabrication and characterisation of reliable G-MEAS still need to be overcome.

## Materials and Methods

### Device fabrication and characterisation

#### Design

The electrodes were designed for a 48 by 48 square mm (mm^2^) borosilicate glass substrate (Diamond Coatings Ltd.) with a thickness of 170 μm (**Fig. 5a**). The device geometry matched the standard head stage of a Multichannel System MEA-2100 Mini Amplifier. The devices consisted of arrays of graphene electrodes placed at the centre of the glass. To connect the electrodes to test instrumentation, gold (Au) leads were fabricated on the same substrate. The Au leads were 15800 μm in length, with 60 Au electrode pads making up the frame of the G-MEA. Each electrode pad was 2200 μm in length and width. The triangular Au feature extending from an electrode pad was the counter electrode used for impedance measurements and the grounding connection (see Final Device in **Fig. 5a**). It was fabricated using Au and had a length of 14400 μm. The triangular feature was 6100 μm in length and 3300 μm in width. There was a 100 μm overlap between the graphene electrodes and Au leads and a 200 μm spacing between each graphene electrode. In the middle of each graphene electrode pad in the centre of the glass, there was a 30 μm diameter hole opening in the SU-8 passivation layer.

**Figure 5.**
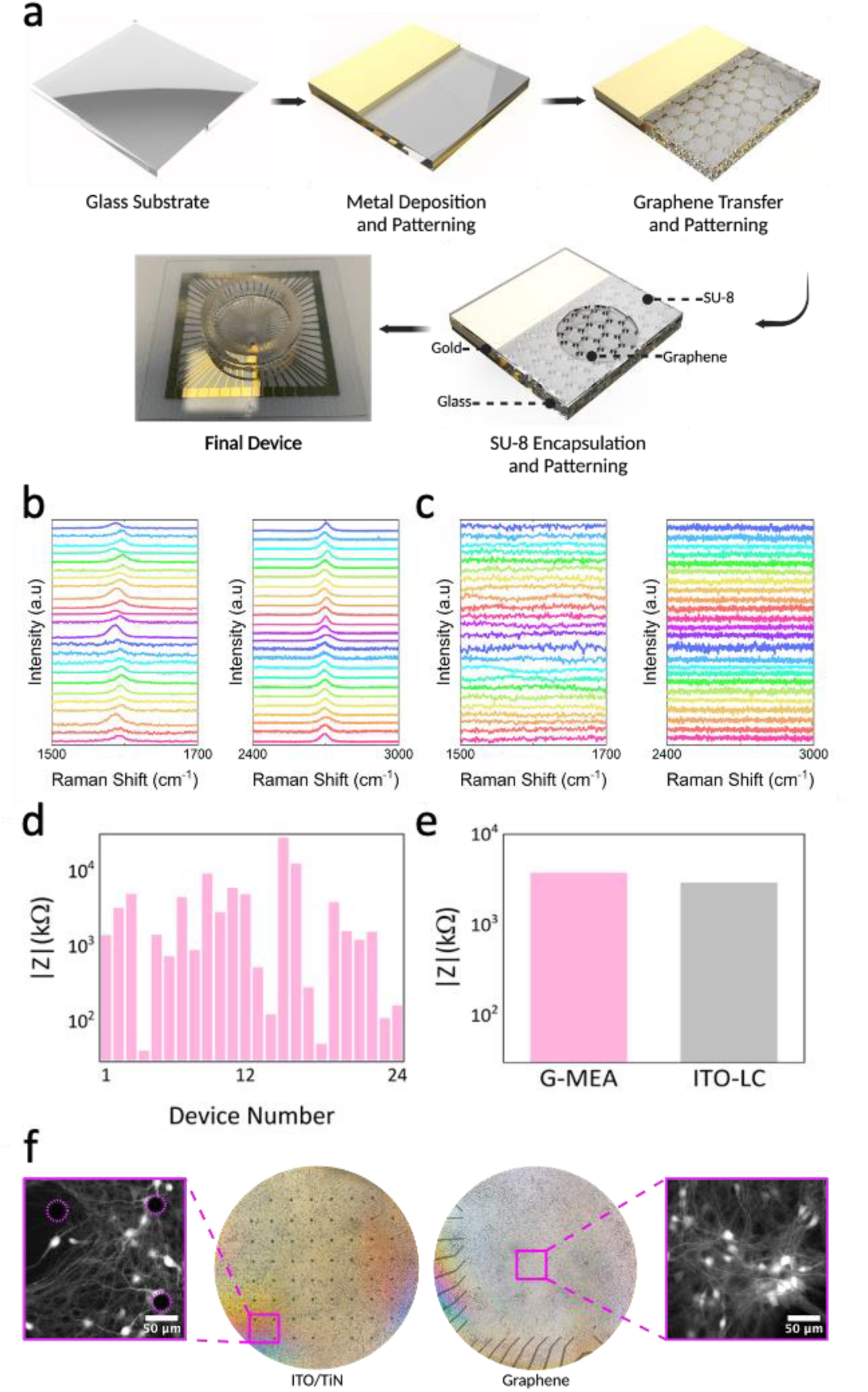
G-MEAs are highly transparent and display a low impedance. (**a**) The main fabrication steps include chromium and gold deposition onto a borosilicate glass coverslip; patterning of gold electrodes using direct laser writing; chemical vapour deposition of graphene and transfer *via* copper etching before patterning graphene electrodes using direct laser writing; and spin-coating of the SU-8 passivation layer onto the device with 30 μm holes being patterned above the graphene electrode pads. (**b**) Comparison of the normalised Raman spectra at 514 nanometres for graphene electrodes across 24 different devices. The characteristic graphene G peaks at ∼1580 cm^-1^ are in the image on the left and 2D peaks at ∼2700 cm^-1^ are in the image on the right. The spectra were obtained to confirm the presence of graphene after the etching and cleaning process. (**c**) Spectra of the areas next to the graphene electrodes after etching to demonstrate the success of the etching and cleaning process across 24 different devices. The image on the left shows the disappearance of the G peaks and the image on the right shows the disappearance of the 2D peaks. (**d**) Comparison of the impedance values across 24 different G-MEAs. The average total impedance at 1 kHz was 3.72 kΩ. (**e**) Comparison of the average total impedances between fabricated G-MEAs and lab-characterised commercial ITO devices (ITO-LC). ITO-LC were characterised using the same technique as for G-MEAs. (**f**) A comparison of the imaging field of view between commercial MEAs and G-MEAs using both bright field and fluorescence microscopy. For the ITO/TiN sample, the black dots indicate the TiN electrodes on top of the connecting ITO tracks. For the graphene sample, the black lines are the gold electrodes which connect to the transparent graphene microelectrodes located inside the square-shaped imaging area (4.84 mm^2^).

#### Fabrication

The glass substrate was initially baked for 20 minutes at 200 ℃ on a hot plate to remove any water molecules on the surface and covered in AZ 5214E photoresist (Merck Performance Materials GmbH) *via* spin coating. The Au leads were patterned by direct laser writing using a 405-nanometre gallium nitride diode laser writer (LW-405B+ Microtech Srl). To avoid unwanted resist exposure caused by light penetrating the glass substrate and getting reflected by the sample holder, the substrate was attached to a silicon substrate using BlackTak^TM^ Light Masking Foil throughout the lithography process. After exposure, the photoresist was developed using an AZ 351B developer (MicroChemicals GmBH) for 51 seconds. Before the metal deposition, the samples were exposed to oxygen plasma (10 Watts for 60 seconds) *via* a Vision 320 Reactive Ion Etcher (Advanced Vacuum AB, Plasma-Therm). Subsequently, 5 nm of chromium was deposited at a rate of 0.5 Å/s, followed by 50 nm of Au at a rate of 0.5 Å/s until it reached five nm, where the rate was increased to 1 Å/s using an electron-beam evaporator (LEV-PVD 200 Pro, Kurt J. Lesker). The chromium was used as an adhesion layer between the Au and the glass substrate. The photoresist was removed *via* lift-off, leaving the sample vertically in 99.9 % analytical-grade acetone (Sigma-Aldrich) for 15 minutes.

Single-layer graphene was produced *via* chemical vapour deposition (CVD) on copper foils^54^. After growth and cooling, poly(methyl methacrylate) 950 A4 (A-Gas Electronic Materials Ltd (Rugby), Kayaku Advanced Materials) was spin-coated onto the graphene on the copper foil, yet not baked. The Cu foil was dissolved in a Cu etchant solution (Sigma-Aldrich), leaving the PMMA+graphene stack floating on the solution. The graphene was cleaned by transferring the stack on a solution of 18.2 MΩ/cm ultrapure water and hydrochloric acid in a 1:10 ratio and further cleaned in new batches of ultrapure water. The PMMA+graphene stack was then transferred onto the Au electrode prefabricated on the glass substrate and left to dry overnight. Subsequently, the samples were submerged in acetone for another night to remove the PMMA layer, releasing the graphene onto the Au leads. Afterwards, the graphene was patterned into the desired electrode shapes using direct laser lithography following the same procedure as described above. The uncovered part of the graphene was etched by oxygen plasma in a reactive ion etcher (Vision 320, Advanced Vacuum AB, Plasma-Therm, 10 W for 3 minutes). The photoresist was then removed by an additional overnight acetone bath. Finally, the device was encapsulated in an SU-8 2000.5 (Kayaku Advanced Materials) passivation layer, and 30 μm hole openings were patterned in the SU-8 by direct laser lithography (375 nm). The holes were precisely aligned with the tip of the graphene microelectrode pads to enable exposure of the graphene surface for sensing. A final hard bake at 180 ℃ was carried out to remove cracks and detoxify the SU-8^55^, before being cooled for 24 hours to reduce stresses in the SU-8. A glass ring with an inner diameter of 19 mm (Diamond Coatings Ltd) was then attached to the G-MEA using a multipurpose silicone-elastomer-based sealant (Dow Corning 732).

#### Characterisation

The presence of graphene microelectrodes on the glass substrate was confirmed using a micro-Raman Spectrometer (Renishaw InVia) at 514 nanometres excitation (**Fig. 5b**). The device was electrically characterised (**Fig. 5c, d**) using a Cascade Summit series semi-automated probe station connected to a 4294A precision impedance analyser (Keysight, Agilent Technologies, model). The measurements were carried out by measuring the total impedance between the counter electrode, the Au ground electrodes on the G-MEAs, and the working electrode, *i.e.*, the graphene microelectrode directly opposite the ground electrode. This was carried out both by measuring the G-MEA without filling the glass cell culture rings of the G-MEAs and immediately after filling the glass cell culture rings with 1300 μL of Dulbecco’s phosphate-buffered saline (Thermo Fisher Scientific). The same electrode and counter electrode were selected for all the G-MEAs measured, and the measurement probes were not moved between devices and measurements. Three commercially obtained ITO MEAs (60tMEA200/30iR-ITO-gr, Multi Channel Systems MCS GmbH) were also characterised in the same manner to compare the impedance values (**Fig. 5e**) and to standardise the impedance measurement methods.

### Cell culture

#### Virus production from HEK cells

Day 1: Plating of 200,000 HEK cells in a plate (Cat. No: 150466, Thermo Fisher Scientific) with antibiotics-containing media. The complete medium for normal cell growth consisted of 90 % DMEM (Sigma-Aldrich), 10 % FBS (Sigma-Aldrich) and 1 % streptomycin (Thermo Fisher Scientific). Day2: HEK cells were incubated in antibiotics-free media for 2 hours before the transduction of plasmid mixtures: prepare a combination of 70 µg of pAAV 2.1 (Rep/Cap), 200 µg of pH GT1-ademo/dF6 helper and 70 µg of pGP-AAV-syn-jGCaMP7b-WPRE (Plasmid Number 104489, Addgene)^56^ and mix with 18 ml OptiMEM (Thermo Fisher Scientific) and 17 ml polyethyleneimine (PEI, Sigma-Aldrich), incubate for 20 minutes at room temperature, and then apply dropwise to the plate. Day 3: change the HEK cell media with antibiotics-containing media but no FBS. Day 4: collect the media from plates to 50 ml tubules (40 ml/tube). Scrape off cells, transfer them to a new tube and spin with cells and media at 2000 rpm for 5 minutes, then transfer media from cells to media tubes. For cells, resuspend cell pellets with AAV lysis buffer up to 1.5 ml total and store at −80°C. Media: for every 40 µl of media add 0.93 g NaCl and 40 µl 40 % polyethylene glycol (PEG, Sigma-Aldrich), and keep at room temperature until NaCl is dissolved. Transfer the media to ice for 2 hours, then spin 10 mins at 6.6 k rpm. Discard media without disrupting the pellets. Resuspend pellets up to 0.5 ml total with AAV lysis buffer.

#### Primary neuron dissection

Hippocampal tissues from 2 days postnatal (P2) rats (Sprague-Dawley rats from Charles River) were dissected and collected in 2 ml Eppendorf tubes containing cold DMEM (Sigma-Aldrich) and maintained on ice. After tissue collection, the cold DMEM was replaced with room temperature DMEM containing 0.1 % Trypsin and 0.05 % DNase (Sigma–Aldrich, UK). The tubes were then incubated in a CO2 incubator set at 37 ℃, 5 % carbon dioxide, and 20 % relative humidity for 20 minutes. Tissues were rinsed four times with 0.05 % DNase in DMEM at room temperature and dissociated into a single-cell suspension by trituration using a 1 ml and then a 200 µL Gilson pipette tip. The cell suspension was centrifuged at 600 rpm for 5 minutes. The supernatant was then discarded, and the pellet was gently resuspended in DMEM containing 10 % FBS. Cell numbers were determined using a haemocytometer.

#### Cell Culture

G-MEAs were filled with poly-L-lysine solution (Sigma-Aldrich) and placed under an ultraviolet lamp in a sterile laminar flow cabinet for one hour. The devices were rinsed with Dulbecco’s Phosphate Buffered Saline (DPBS) before 1300 microlitres (μL) of the neurobasal medium was introduced. Neurobasal media was used for the maintenance and maturation of the cells by improving cell survival through the provision of supplements. Neurobasal media contained 2 % B27 and 0.25 % Glutamax (all from Thermo Fisher Scientific). The devices were placed in an incubator set at 37 ℃, 5 % carbon dioxide, and 20 % relative humidity to warm up before plating primary P2 hippocampal neurons. Rat primary neurons were isolated as mentioned above. 255,000 primary hippocampal cells were plated directly into the middle of the device, where the graphene microelectrodes sat. 200 μL of media was taken out and replaced by 300 μL of warmed-up neurobasal medium every other day to maintain the cell culture in an incubator set at 37 ℃, 5 % carbon dioxide, and 20% relative humidity. On DIV 4, the hippocampal neurons were transfected using GCaMP7b, using an adeno-associated virus vector pGP-AAV-syn-jGCaMP7b-WPRE (see production above). At the start of DIV 24, selected neurons were imaged in each quadrant of the device and simultaneous electrophysiology recordings of the selected neurons were obtained. At the end of DIV 24 (Day 1), after the baseline imaging and electrophysiology recordings were obtained, 10 μM of the drug U18666A (662015, Sigma-Aldrich) were introduced by triturating the drug and the device media three times gently. This was carried out every day until DIV 27 (Day 4).

### Imaging and electrophysiology

#### Widefield Imaging

Calcium imaging was carried out using a custom-built automated wide-field microscope (IX83, Olympus), with an sCMOS camera (Zyla 5.5, Andor), and a four-wavelength high-power light emitting diode light source (LED4D067, Thorlabs). The software Micro-Manager^57^ was used to control the system.

#### Simultaneous Electrophysiology and Imaging

The MEA head stage was mounted onto the microscope imaging stage. A lens extender was used to increase the height of the microscope lens so that the lens could pass through the base of the head stage and form contact with the bottom of the G-MEA. A stage-top heater (OKOLab, Ottaviano, Italy) was used to ensure the temperature of the media in the G-MEA did not fluctuate. The heater was set at 37 ℃, 5 % carbon dioxide, and 20 % relative humidity. On DIV 23, sequential images of the transparent FOV were captured and stitched together. The stitched image was overlayed with an image of the graphene electrodes, to create a digital map of the whole device. The image of the device was then split into four quadrants. Three quadrants were used for widefield imaging while the final quadrant was reserved for structured illumination microscopy. This was to minimise the effects of phototoxicity on the cells. Neurons were then selected in each quadrant and marked on this map for simultaneous imaging and electrophysiology over the next four days (DIV 24 - 27). On the second (DIV 25) and fourth day (DIV 27), the neurons were imaged, both using widefield microscopy and super-resolution microscopy, and electrophysiological recordings were obtained. Electrophysiology and microscopy measurements were taken simultaneously. The neurons were imaged over 20,000 frames with a zero-millisecond interval, at an exposure time of 10 milliseconds using a 3×3 binning, and the overall imaging window was around 250 sec. This resulted in every graphene MEA having 30 minutes of electrophysiological recordings and 15 minutes of calcium imaging recordings each day. Calcium fluorescent intensity of the region of interest was extracted and analysed using a custom Python code. The electrophysiology traces and calcium imaging traces were manually aligned using a python script to record the time difference between the initiation of the recordings for the MEA head stage and the image acquisition, which was then used as the time difference value for trace alignment. Based on the script initiation times and CPU testing, there was an unaccounted delay of less than one second, estimated to around 0.6 seconds.

Tiling of images to generate the whole FOV was required due to the high transparency (**Fig. 5e**). Tiling was conducted using a custom-built algorithm using MATLAB. To measure neuron density, we first tuned the brightness of all the widefield images to make them consistent in background fluorescence intensity by Fiji. We then binarised all the frames and measured the density of fluorescence in each of them by Fiji and analysed the extracted data of fluorescence intensity by a custom-built algorithm and GraphPad Prism 9.5.1. All custom designed scripts can be found at https://github.com/MariusBrockhoff/GrapheneMEASpikeSortImaging.

#### Structured Illumination Microscopy

SIM imaging was performed using a custom-built imaging system based on an Olympus IX71 microscope stage, as previously described^48^. Fluorescence excitation of the samples was achieved using a laser emitting at a wavelength of 488 nanometers (iBEAM-SMART-488, Toptica Photonics). The laser beam was expanded to fill the display of a ferroelectric binary Spatial Light Modulator (SLM) (SXGA-3DM, Forth Dimension Displays) to pattern the light with a grating structure. The polarization of the light was controlled with a Pockels cell (M350-80-01, Conoptics). A 60x/1.2 numerical aperture (NA) water immersion lens (UPLSAPO 60XW, Olympus) focused the structured illumination pattern onto the sample. The fluorescence emission from the samples was captured and projected onto an sCMOS camera (C11440, Hamamatsu). The maximum laser intensity at the sample was 20 W/cm^2^. Raw images were acquired with the HCImage software (Hamamatsu) to record image data to disk and a custom LabView program (available upon request) to synchronise the acquisition hardware. Multicolour images were registered by characterising channel displacement using a matrix generated with TetraSpeck beads (Life Technologies) and imaged in the same experiment as the cells.

Resolution-enhanced images were reconstructed from the raw SIM data with LAG SIM, a custom plugin for Fiji/ImageJ available in the Fiji Updater. LAG SIM provides an interface to the Java functions provided by fairSIM^58^. LAG SIM allows users of our custom-built microscope to quickly iterate through various algorithm input parameters to reproduce SIM images with minimal artefacts; integration with Squirrel^59^ provides a numerical assessment of such reconstruction artefacts.

### Data analysis and machine learning

#### Calcium Imaging

Fluorescence calcium signals of spatial areas, *e.g.*, single neurons, of the acquired images were extracted with ImageJ (mean intensity of ROI). Subsequently, these traces were corrected for bleaching by fitting a polynomial decreasing function to the raw data. From the corrected traces, fluorescence calcium spikes were identified, and the respective amplitude and spike time were extracted by finding the local maxima of the trace. The calcium spikes were analysed to extract parameters such as calcium spike frequency and synchronicity^60^ between single neurons in the same FOV.

#### MEA Electrophysiology

Raw MEA recording data were first bandpass filtered (300–-3000 Hz), followed by threshold-based detection of spikes^25^. Extracted spikes were filtered for artefacts by applying machine learning (ML)-based spike sorting on all available spikes of each experiment, manually deciding which spike classes (forcing, *e.g.,* sorting into 20+ classes) represent noise or artefacts. The remaining true spikes were analysed to extract frequently used parameters such as spike rate and mutual information between electrodes.

#### ML for Spike Sorting

The general workflow of spike sorting usually consists of 5 operations: Filtering, spike detection, data pre-processing, feature extraction, and, finally, clustering. While ML has been applied to multiple of those steps^29^, we focused on optimising a combined feature extraction and clustering algorithm. We employed and translated several established deep clustering approaches to the spike sorting task, namely Deep Embedded Clustering^31^ (DEC) and Improved Deep Embedded Clustering^32^ (IDEC). DEC simultaneously learns feature representations and cluster assignments by learning a mapping from the data space to a lower-dimensional feature space in which it iteratively optimises a clustering objective. IDEC is based on integrating the clustering loss and autoencoder’s reconstruction loss: IDEC can jointly optimise cluster label assignment and learn features that are suitable for clustering with local structure preservation. All methods have been implemented in Python, based on TensorFlow^61^/Keras^62^, and can be found at https://github.com/MariusBrockhoff/GrapheneMEASpikeSortImaging.

## Supplementary Materials

### Large-scale synthetic dataset for development and benchmarking of the spike sorting algorithm

The presented datasets aim to provide an extensive, ground-truth basis for the evaluation and benchmarking of spike-sorting algorithms. Unfortunately, currently available datasets are too simple and small to realistically allow to capture the recent, mostly machine-learning-driven, advances in the field or to reflect the magnitude of spikes recorded by the latest generation of MEAs. The datasets contain simulated spike shape recordings generated *via* NeuroCube^33^. To create the recordings, the default settings of NeuroCube have been used (single electrode, 300.000 neurons/mm^3^, 7 % rate of active neurons, exponential firing rates). Recordings were simulated with a sampling frequency of 20 KHz. For each recording, five neurons (maximum available) have been manually placed around a single recording electrode. The relative distance to the electrode has been randomly sampled between 0 and 1 (step size 0.01) and firing rates have been randomly sampled between 15 and 35 Hz. Alongside the datasets, we provide the created Cube files that can be loaded into the NeuroCube software in order to simulate the same conditions and neurons that have been used here.

**SI Table 1:**
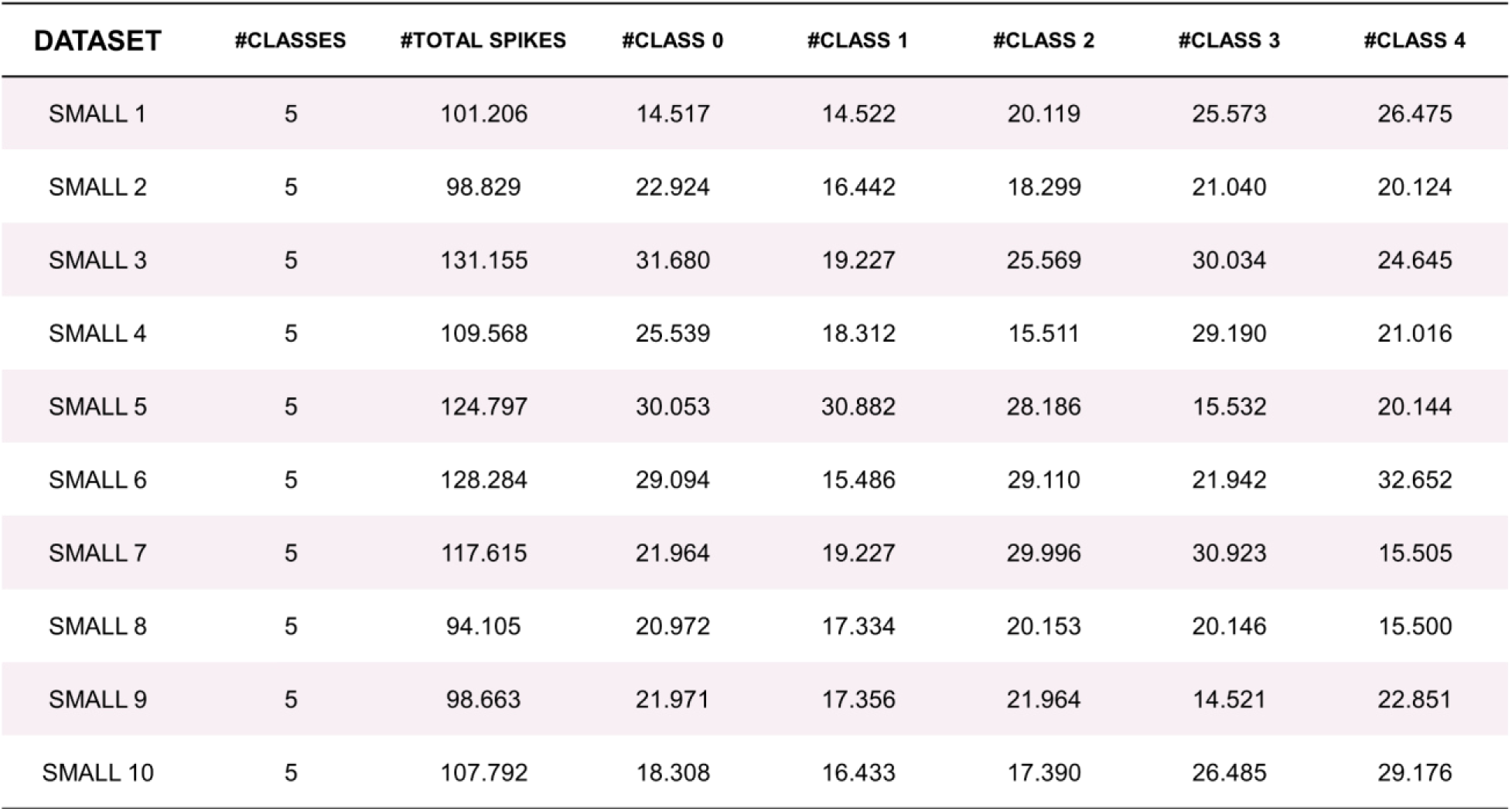
Overview number of spikes per class for the *Small* dataset.

Data is stored as Python pickle files. Each pickle file can be loaded with a simple pickle.load() command. Stored is an array of dimensions Number_of_spikes x 66. The first column carries the ground-truth spike class for each spike (integer). The second column contains the spike time of the simulated spike (in ms). Finally, columns 3 to 66 contain the respective spike shape (64 data points).

The datasets are organised by size and complexity. The group of sets called *Small* are 10 sets of spike recordings, each including about 100,000 spike shape recordings of 5 classes (source neurons). The spike shapes and firing rates present in each set have been chosen randomly. Details on spike shapes as well as the number of recordings of each neuron can be found in Supplementary Table 1.

The datasets called *Large* are 10 sets of spike recordings, each including about 1,100,000 spike shape recordings of 5 classes (source neurons). The spike shapes and firing rates present in each set have been chosen randomly. Details on spike shapes as well as the number of recordings of each neuron can be found in SI Table 2.

**SI Table 2:**
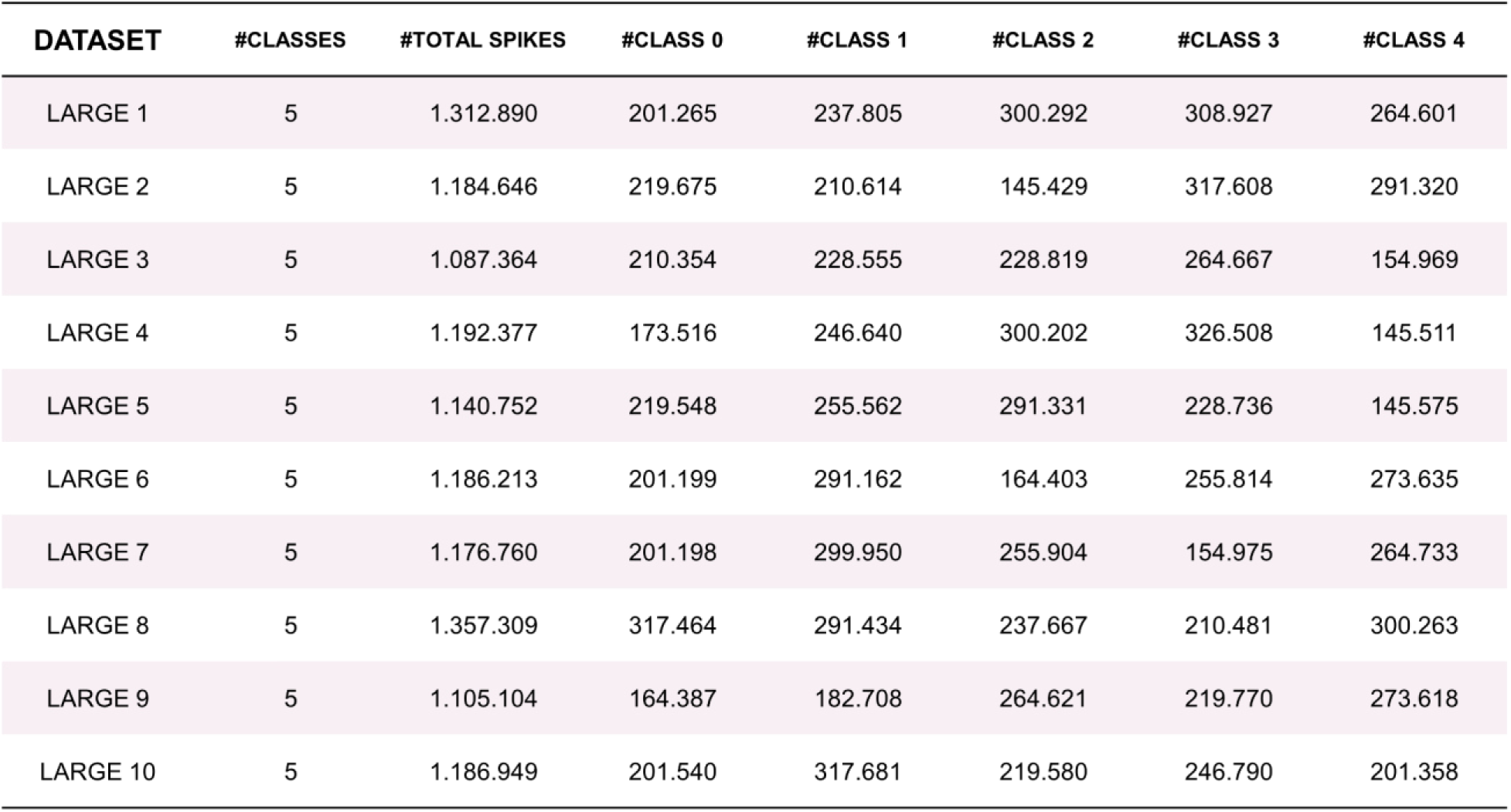
Overview number of spikes per class for the *Large* dataset.

### Benchmarking deep clustering algorithms on large-scale synthetic dataset

**SI Table 3:**
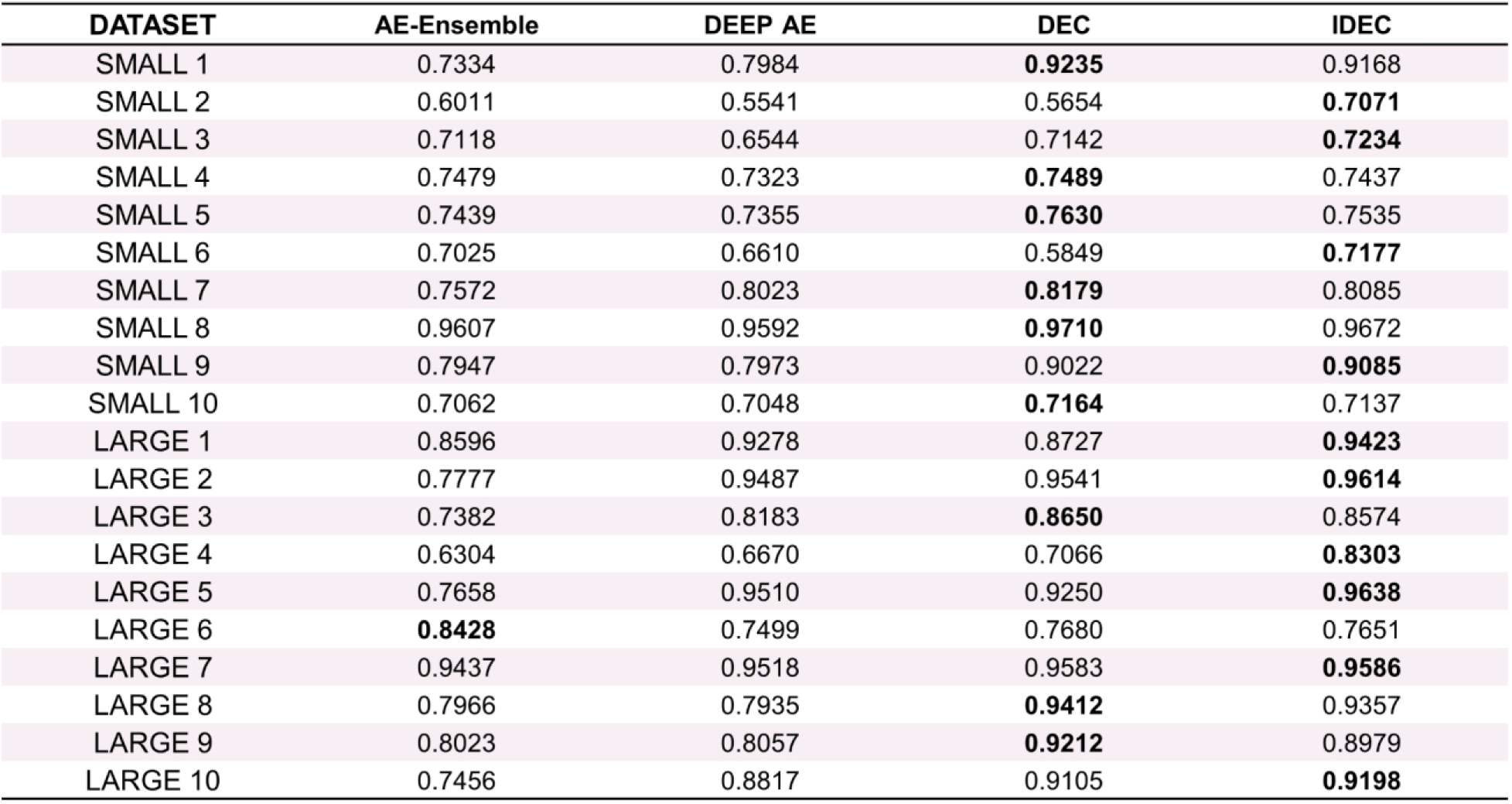
Benchmarking performance (accuracy) of several ML-based spike sorting approaches. Given is the best result of 5 repeated runs. To enable a fair comparison, we compare accuracies obtained *via* K-means++^63^ clustering with the correctly given number of clusters. Best performing model per dataset is marked in bold.

**Supplementary Figure 1.**
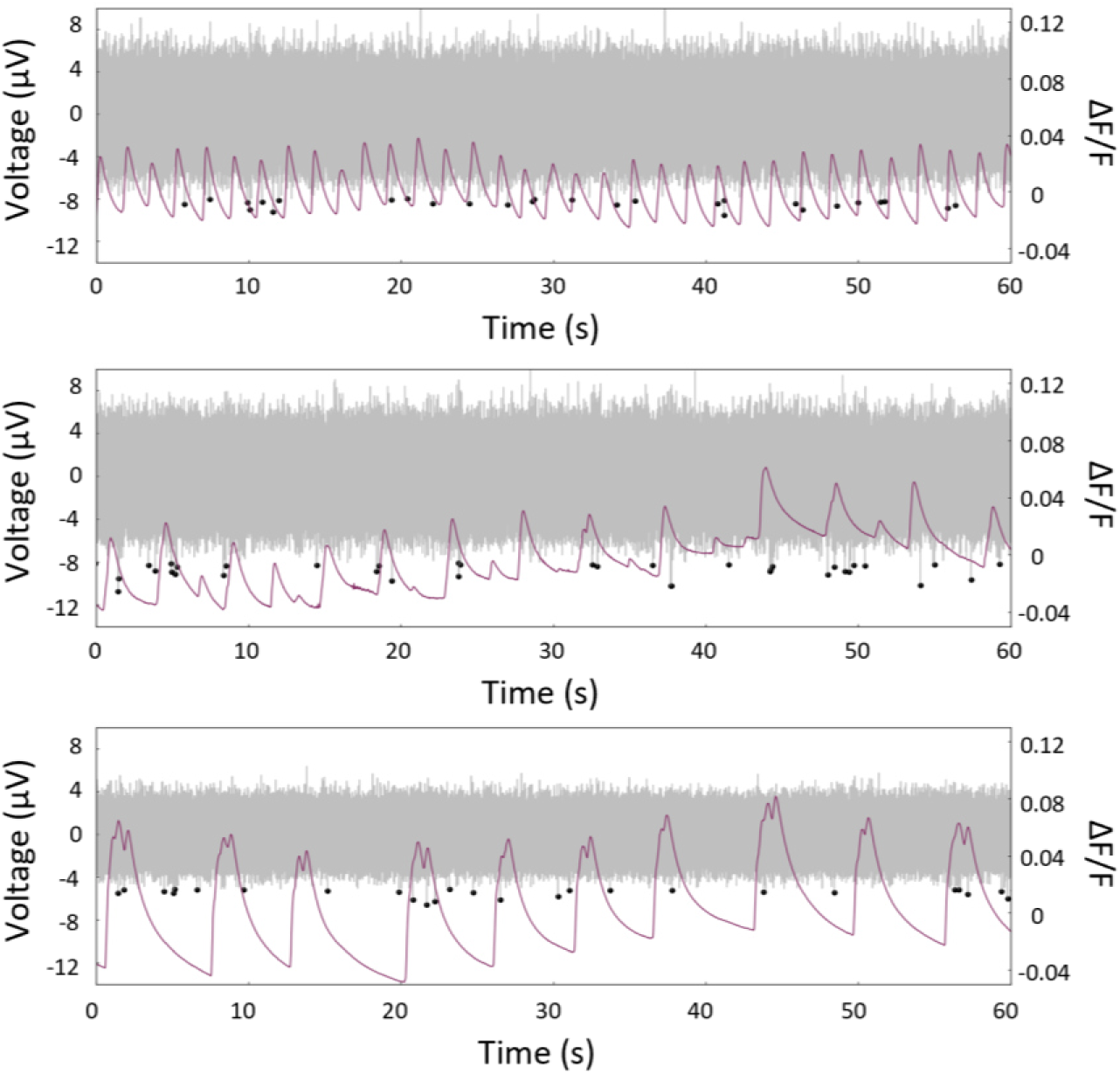
Correlated local electrophysiological and calcium activity recordings from different locations on G-MEAs highlight variations in signals including reduced correlation between events of the signal types as well as differences in firing rates.

**Supplementary Figure 2.**
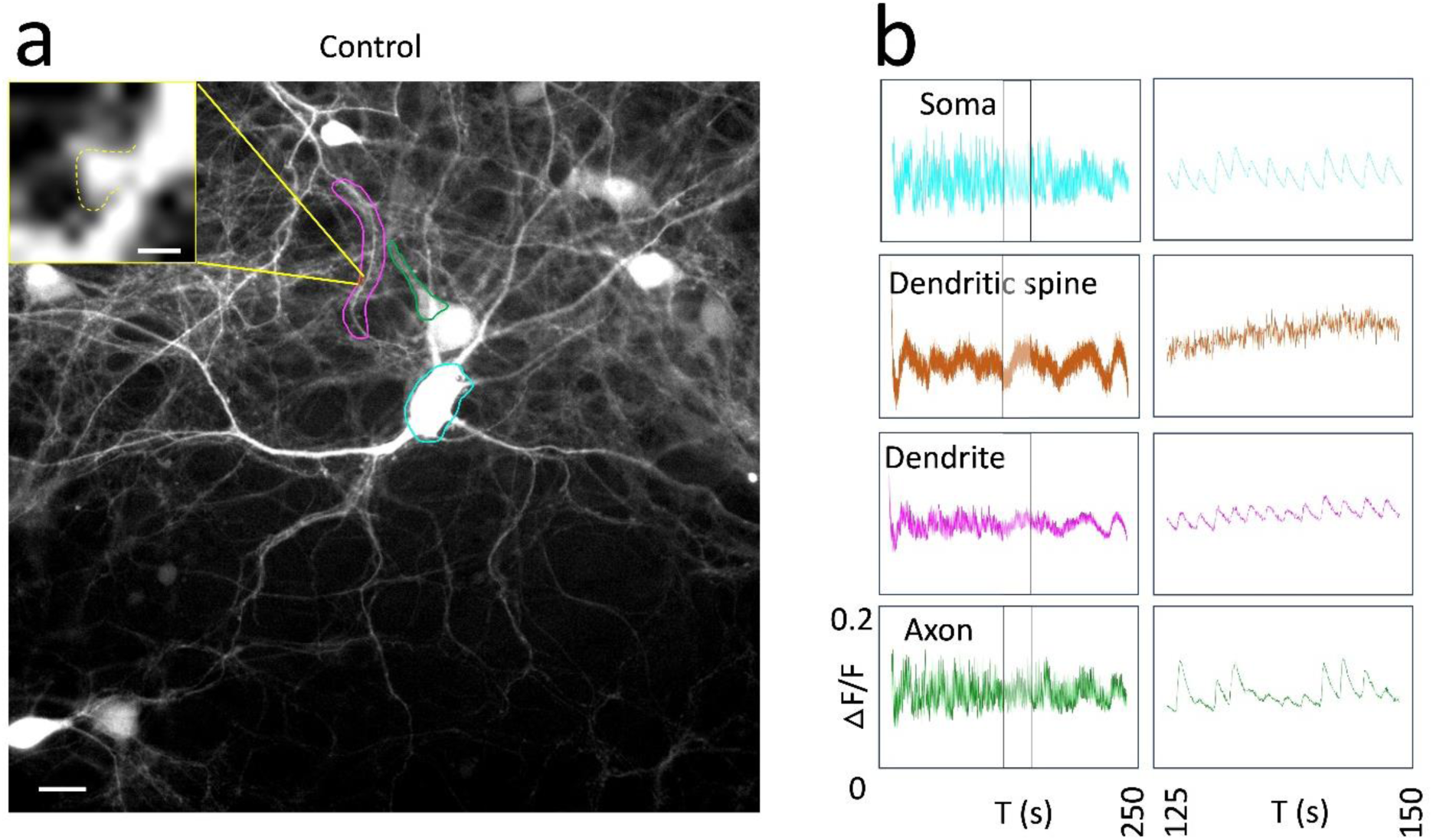
(**a**) The structure of a single neuron in a single FOV. Sub-neuronal structures highlighted in the figure are a dendrite (magenta), an axon (green), a soma (cyan), and a dendritic spine (brown). The enlarged boxed region highlights the dendritic spine analysed in b. Scale bars: 20 µm, 1 µm for enlarged FOVs. (**b**) Changes in the amplitude of the calcium sensor fluorescence extracted from the neuronal structures are depicted over a time window of 250 s. Y-axis: ΔF/F with a scale of 0 to 0.6 for all the plots. The small time-window (25 s, right side) is indicated by black rectangles in the full-time window panel (left side). Sub-plots have the same y-axis and scales.

## Acronyms

G-MEAs: Graphene microelectrode arrays
NPC: Niemann-Pick disease type C
MEAs: Microelectrode arrays
FOV: Field of view
ITO: Indium tin oxide
ML: Machine Learning
SIM: Structured illumination microscopy
DEC: Deep Embedding for Clustering
IDEC: Improved Deep Embedding for Clustering
AE-Ensemble: Autoencoder-Ensemble
Deep AE: Standard deep autoencoder
DIV: Day *in vitro*
AAV: Adeno-associated viruses
4D-SIM: Four-dimensional structured illumination microscopy
3D: Three-dimensional
FLIM: Fluorescence lifetime imaging microscopy
FRET: Fluorescence resonance energy transfer

## Units

kilo Ohms (kΩ)

micrometres (μm)

reciprocal centimetre (cm^-1^)

millimetres squared (mm^2^)

kilo Hertz (kHz)

